# Associations between abstract working memory abilities and brain activity underlying long-term recognition of auditory sequences

**DOI:** 10.1101/2022.05.19.492607

**Authors:** G. Fernández Rubio, F. Carlomagno, P. Vuust, M. L. Kringelbach, L. Bonetti

## Abstract

Memory is a complex cognitive process comprised by several subsystems, namely short- and long-term memory and working memory (WM). Previous research has shown that adequate interaction between subsystems is crucial for successful memory processes such as encoding, storage and manipulation of information. However, few studies have investigated the relationship between different subsystems at the behavioral and neural levels. Thus, here we assessed the relationship between individual WM abilities and brain activity underlying the recognition of previously memorized auditory sequences.

First, recognition of previously memorized versus novel auditory sequences was associated with a widespread network of brain areas comprising the cingulate gyrus, hippocampus, insula, inferior temporal cortex, frontal operculum, and orbitofrontal cortex.

Second, we observed positive correlations between brain activity underlying auditory sequence recognition and WM. We showed a sustained positive correlation in the medial cingulate gyrus, a brain area which was widely involved in the auditory sequence recognition. Remarkably, we also observed positive correlations in the inferior temporal, temporal-fusiform, and postcentral gyri, brain areas which were not strongly associated to auditory sequence recognition.

In conclusion, we discovered positive correlations between WM abilities and brain activity underlying long-term recognition of auditory sequences, providing new evidence on the relationship between memory subsystems. Furthermore, we showed that high WM performers recruited a larger brain network including areas associated to visual processing (i.e., inferior temporal, temporal-fusiform and postcentral gyri) for successful auditory memory recognition.

**Significance statement:** Memory is a complex cognitive process dependent on the successful interaction between its multiple subsystems. Here, we assessed the relationship between individual WM abilities and brain activity underlying the recognition of previously memorized auditory sequences.

We observed positive correlations between brain activity underlying auditory sequence recognition and WM, especially in the medial cingulate gyrus, inferior temporal, temporal-fusiform and postcentral gyri. In this study, we provided new evidence on the relationship between two memory subsystems: WM and long-term auditory recognition. Moreover, we showed that, to successfully complete memory recognition tasks, high WM performers recruited a larger brain network which comprised brain areas mainly associated to visual processing, such as inferior temporal, temporal-fusiform and postcentral gyri.

## Introduction

Memory is a fundamental cognitive process that is widely regarded as a multisystem function ^1^ relying on a widespread network of brain areas such as the medial temporal lobe ^2, 3^, prefrontal cortex ^4^, and basal ganglia ^5^. Broadly, the memory subsystems encode, store, and retrieve past memories (long-term memory), temporarily store sensory information (short-term memory), and maintain and manipulate data (working memory) ^1, 6, 7^. These subsystems operate simultaneously and in parallel ^8^, giving rise to efficient memory functioning that is essential for many daily activities.

Working memory (WM) capacity allows to briefly store and manipulate information and is involved in decision-making and executive processes ^9-11^. Among the several theories of WM, Baddeley and Hitch’s ^12^ multicomponent model has become highly influential. According to this theory and its subsequent revisions, WM is comprised by four components: (1) the phonological loop, which is involved in verbal WM, (2) the visuospatial sketchpad, for visuospatial WM, (3) the central executive, or the attentional control system, and (4) the episodic buffer, for storing information ^10, 12-14^. Frequently, WM paradigms request individuals to retain sensory information and perform some operation or manipulation on it, as in the case of the *N*-back ^15^ and digit span ^16^ tasks.

Neuroimaging studies have highlighted the role of cortical brain areas, such as the prefrontal, parietal and cingulate cortices, and subcortical areas including the midbrain and cerebellum in WM processes, as reported in a review by Chai et al. ^17^. Evidence comes mainly from studies using visual stimuli, providing a valuable but incomplete picture of the neuroanatomy of WM. However, recent studies on auditory WM processing have uncovered the role of the primary auditory cortex and high-order structures such as the hippocampus for this cognitive function. For example, Kumar and colleagues ^18^ demonstrated that the activity and connectivity of the primary auditory cortex, hippocampus and inferior frontal gyrus are associated with the maintenance of single sounds’ series. Additionally, theta oscillations and phase locking in the dorsal stream predicts performance in a maintenance and manipulation auditory task ^19^. Related to the present study, Bonetti et al. ^20^ showed a positive correlation between WM capacity and brain activity underlying an auditory mismatch-negativity (MMN) task. The authors found that participants with higher WM scores showed enhanced MMN responses in frontal regions, but not in temporal areas. Notably, this investigation evidenced the relationship between auditory short-term and working memory.

Long-term memory refers to the ability to recall information that has been encoded and stored in the past ^7, 21^. Research on this cognitive function has emphasized the distinct features of several types of long-term memory, namely episodic, semantic, and procedural memory ^22, 23^. These are classified according to the kind of information they hold (e.g., personal experiences in the case of episodic memory, factual knowledge for semantic memory) ^24, 25^ and how this information is encoded (e.g., skill acquisition in procedural memory) ^26^.

The neural underpinnings of long-term memory rest primarily upon medial temporal lobe structures (hippocampus, entorhinal, perihinal and parahippocampal cortices) ^2, 21^ and interact with the prefrontal cortex for successful memory retrieval ^27^. Moreover, consolidation, the process of transforming temporary information into long-lasting memories and a central aspect of long-term memory, is achieved through the interactions between the hippocampus and neocortex ^28, 29^. Converging evidence suggests that, in the case of auditory long-term memory, the primary auditory cortex also supports the storage of information ^30^.

Although previous investigations have mainly examined the neuroanatomical bases of the memory subsystems in isolation, a few studies have looked into the associations between them. For instance, Henson and Gagnepain ^31^ highlighted the interaction between different memory subsystems, both in terms of behavior and neural substrate. They focused especially on episodic, semantic, and modality-specific perceptual subsystems, claiming that their successful interaction is crucial for performing memory tasks. Similarly, Poldrack and colleagues ^32^ demonstrated the interaction and competition between memory subsystems during classification learning in humans. Specifically, they observed that the basal ganglia and medial temporal lobe were differently engaged depending on the emphasis on declarative or non-declarative memory and showed that the interaction between these structures was necessary to perform the task. In a review focusing on pharmacological and neurochemical studies, Gold ^33^ proposed that the release of acetylcholine in different memory subsystems showed extensive interactions between them, which could be cooperative or competitive. He concluded that different memory and neural systems tended to interact extensively, even when described as relatively independent. Finally, White and McDonald ^34^ described a theory of multiple parallel memory subsystems in the rat brain localized in the hippocampus, caudate-putamen, and amygdala. The authors claimed that all subsystems had access to the same information during learning, but that each subsystem represented a different relationship between the information features. In their view, these memory subsystems interacted by simultaneous parallel influence on behavioral output and by directly affecting each other in a cooperative or competitive manner. Overall, these investigations have yielded considerable insights into the relationships between memory subsystems, but we still lack information on the brain correlates underlying these interactions.

Thus, in our study we aimed to investigate the relationship between two of the most important memory subsystems, WM and long-term memory, emphasizing their interdependence. To this end, we correlated the scores from a widely used WM measure with the neural activity underlying tone-by-tone recognition of previously memorized sequences from three different musical pieces. We hypothesized to observe stronger brain activity underlying auditory sequence recognition in individuals with greater WM abilities, especially in brain structures that have been previously associated to memory processes, such as the prefrontal cortex and hippocampus. Additionally, we expected WM capacity to be positively correlated with behavioral responses in the auditory recognition task.

## Results

### Experimental design

Participants performed an old/new auditory recognition task ^35-37^. During the encoding phase, participants listened to three musical pieces and were instructed to memorize them as much as possible. In the recognition phase, short musical sequences selected from the pieces (i.e., memorized musical sequences) and novel musical sequences were presented. For each of the sequences, participants stated whether they memorized or novel. Their brain activity was recorded using magnetoencephalography (MEG) during the recognition task. Structural magnetic resonance imaging (MRI) images were collected for each participant and combined with the MEG data to reconstruct the sources using a beamforming approach, which generated the signal that recorded over the MEG channels. Finally, participants’ WM abilities were measured using the Digit Span and Arithmetic subtests from the Wechsler Adult Intelligence Scale (WAIS-IV) ^38^. **Figure 1** shows a graphical depiction of the experimental design and analysis pipeline.

**Figure 1.**
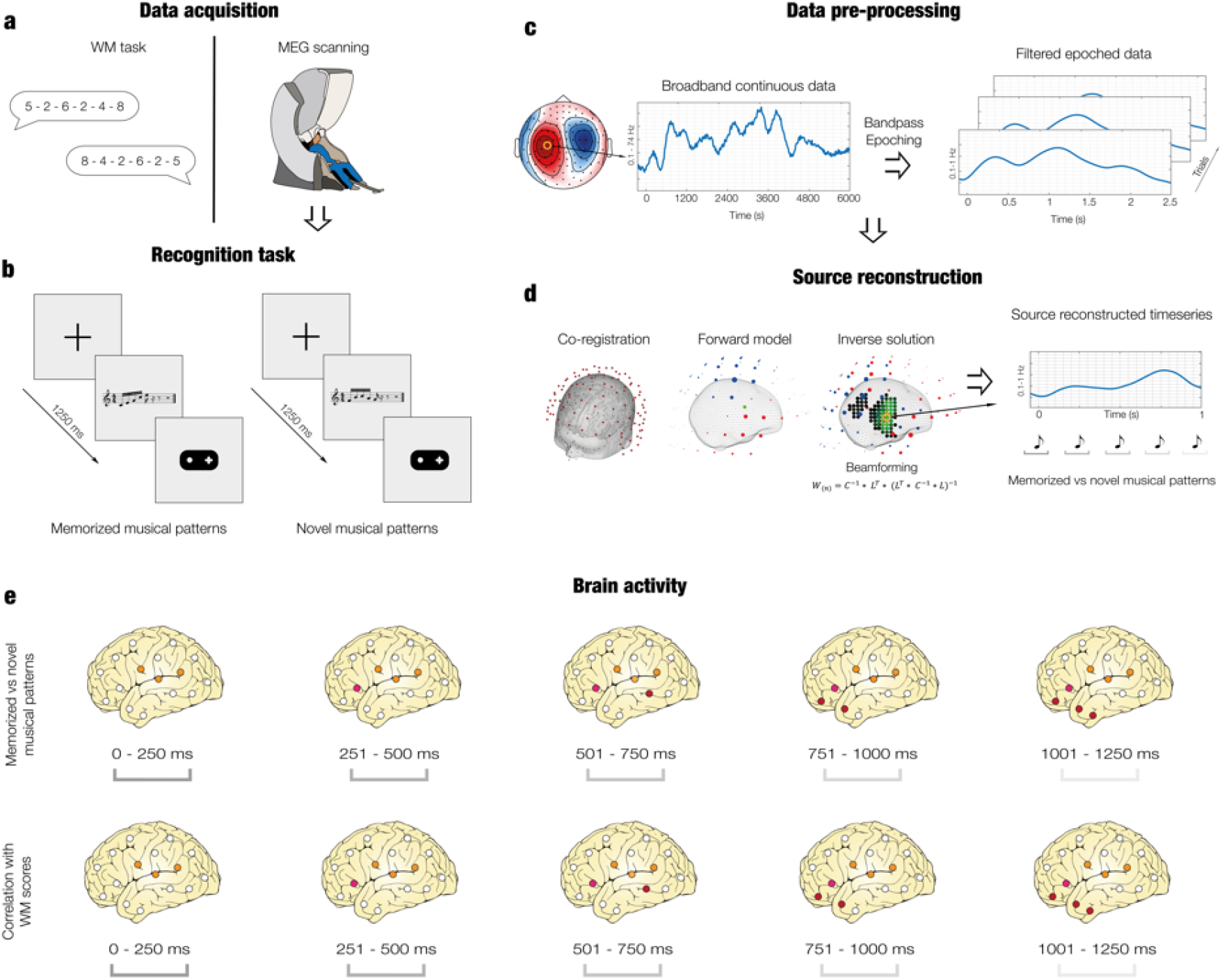
Experimental stimuli and design, and data analyses overview. **a** – The data acquisition comprised two parts: a working memory (WM) task completed outside the scanner and an old/new auditory recognition task that was carried out during MEG recording. **b** – Illustration of the old/new auditory recognition task performed in the MEG scanner. After listening to three full musical pieces, participants were presented with melodic excerpts that were extracted from the pieces they previously learned or with new melodies, and were asked to state whether each melody was memorized or novel using a joystick. **c** – The broadband continuous neural data was preprocessed, bandpass filtered (0.1-1Hz), and epoched. **d** – Source reconstruction analyses were performed to isolate the contribution of each brain source to the neural activity recorded by the MEG sensors. Preprocessed MEG and MRI data were co-registered. After that, a forward model was computed, and the inverse solution was estimated using a beamforming approach. **e** – Contrasts between memorized and novel auditory sequences were calculated for each musical tone (top row). Pearsons’ correlations between WM scores and brain activity underlying recognition of memorized versus novel auditory sequences were computed (bottom row).

### Brain activity underlying recognition of previously memorized versus novel musical sequences

Before evaluating the relationship between WM abilities and brain activity underlying musical sequence recognition, which was the main aim of the current work, we wished to replicate the established finding ^35-37^ that recognition of previously memorized versus novel auditory sequences is associated to a stronger activation in a widespread network of brain areas.

First, we sub-averaged the brain data in five time-windows, corresponding to the duration of the five tones of the musical sequences (0 – 250 ms, 251 – 500 ms, 501 – 750 ms, 751 – 1000 ms, 1001 – 1250 ms). Second, independently for the five time-windows, we computed one t-test for each brain source, contrasting the brain activity underlying recognition of previously memorized versus novel musical sequences. Third, we corrected for multiple comparisons by using cluster-based Monte-Carlo simulations (MCS).

Significant clusters of activity (*p* < .001) were located across a number of brain voxels (*k*) for each tone of the musical sequences. As expected, the main clusters were observed for the third (*k* = 284), fourth (*k* = 390), and fifth tones (*k* = 125). The strongest differences between the two conditions were localized in the middle cingulate gyrus, precuneus, insula, hippocampal regions, orbitofrontal cortex, and frontal operculum.

Detailed statistics and information for each voxel forming the significant clusters are reported in **Table ST1**, while a graphical depiction of the results is illustrated in **Figure 2a**.

**Figure 2.**
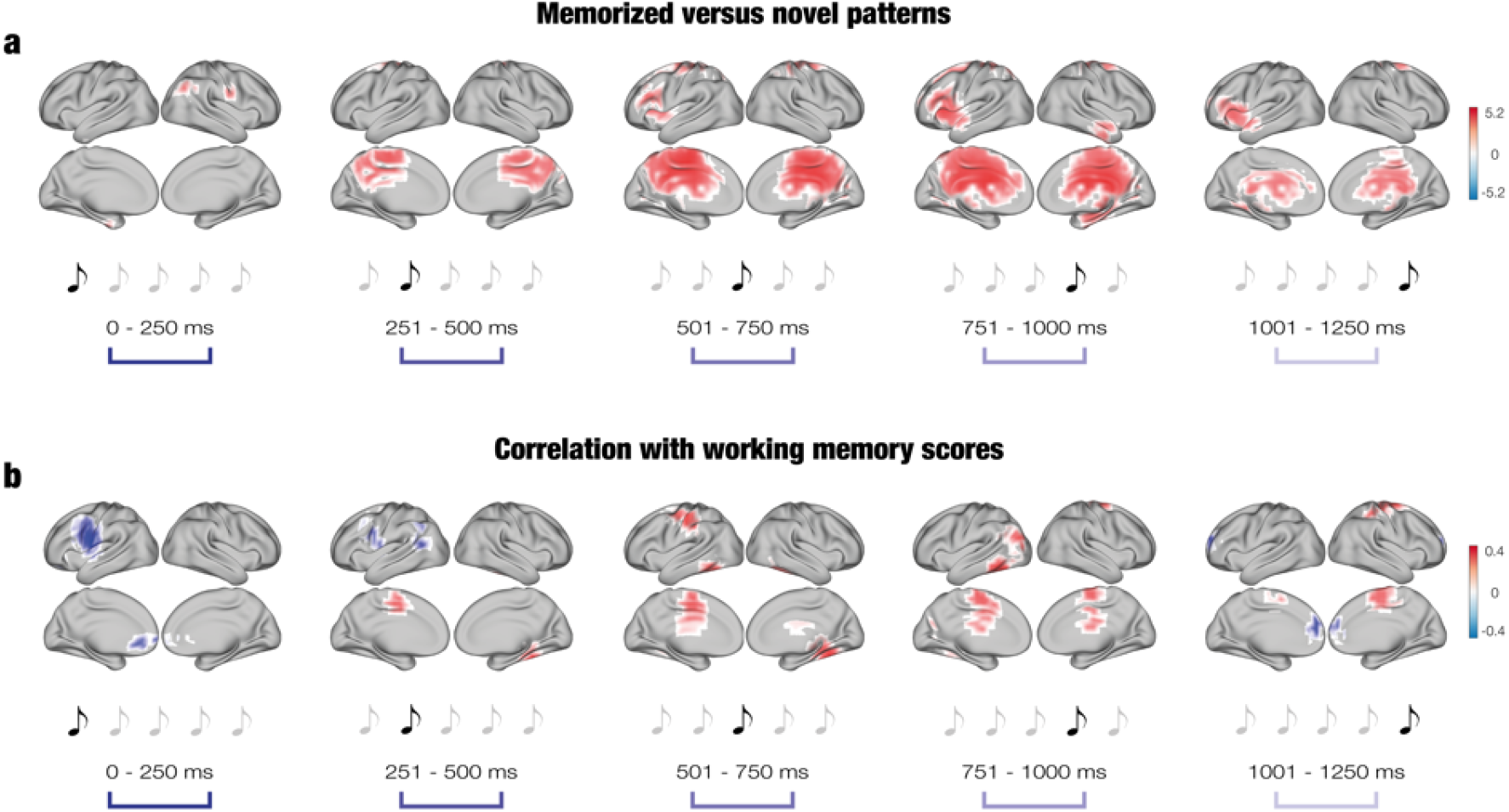
Brain activity underlying the recognition of auditory sequences and correlation with WM scores. **a** – Significant brain activity underlying recognition of the musical sequences. The activity is depicted in brain templates in five subsequent time windows corresponding to the duration of each musical tone forming the sequences (as illustrated by the sketched musical tones above the time windows). The colorbar shows the t-values resulting from the contrast between memorized and novel auditory sequences. **b** – Significant Pearson’s correlations between the brain activity underlying recognition of the sequences and WM scores. The correlations are depicted in brain templates in five subsequent time windows corresponding to the duration of each musical tone forming the sequences (as illustrated by the sketched musical tones above the time windows). The colorbar shows the Pearson’s correlation coefficient obtained by correlating the brain activity underlying recognition of the previously memorized versus novel auditory sequences with the WM scores.

### WM abilities and brain activity underlying musical sequence recognition

The main aim of the study was to establish whether there was a significant relationship between WM abilities and brain activity underlying tone-by-tone recognition of musical sequences.

Before computing neural data analyses, we calculated a Pearson’s correlation between the number of correctly recognized auditory sequences in the MEG task and the individual WM scores. The analysis returned a non-significant result (*rho* = .16, *p* = .18).

To address our experimental question, we computed Pearson’s correlations between participants’ WM scores and each of the reconstructed brain sources. We corrected for multiple comparison using cluster-based Monte-Carlo simulations (MCS). This procedure was computed independently for five time-windows, corresponding to the duration of the five tones of the musical sequences (0 – 250 ms, 251 – 500 ms, 501 – 750 ms, 751 – 1000 ms, 1001 – 1250 ms; see Methods for details).

Significant clusters of activity (*p* < .05) were located in different brain regions and depicted an overall positive correlation between WM abilities and brain activity underlying recognition of memorized musical sequences. This difference returned consistent clusters in the middle cingulate gyrus, inferior temporal cortex, fusiform-temporal cortex, para-hippocampal gyrus, and temporal-occipital fusiform cortex, especially for the third (*k* = 83) and fourth (*k* = 83) tones of the musical sequences.

Detailed statistics and information for each voxel forming the significant clusters are reported in **Table ST2**, while a graphical depiction of the results is illustrated in **Figure 2b**.

## Discussion

In this study, we assessed the relationship between individual WM abilities and brain activity underlying long-term recognition of auditory sequences.

First, we identified the brain activity associated to recognition of previously memorized versus novel auditory sequences. This analysis revealed a widespread network of brain areas involved in the recognition process including the cingulate gyrus, hippocampus, insula, inferior temporal cortex, frontal operculum, and orbitofrontal cortex. Remarkably, the cingulate gyrus (especially the posterior part) was significantly more active for memorized than for novel sequences by the second tone of the sequence. Moreover, this region was strongly active during processing of the rest of the sequence, although its activity decreased in the last tone. Conversely, the insula, inferior temporal cortex and hippocampal areas were mainly active during the third, fourth and fifth tones of the auditory sequence.

Second, we correlated the brain activity underlying recognition of memorized versus novel sequences with the participants’ WM scores. In general, we observed positive correlations between brain activity and WM capacity. The analyses returned a sustained positive correlation in the medial cingulate gyrus, a brain region strongly involved in the auditory sequence recognition. Notably, we also observed positive correlations in the inferior temporal, temporal-fusiform and postcentral gyri. These brain areas were not strongly associated to auditory sequence recognition and suggest that high WM performers may recruit a larger brain network to successfully complete memory recognition tasks.

Our results on the whole-brain mechanisms for auditory recognition are coherent with previous studies that employed the same paradigm. For instance, using part of the current dataset, Bonetti et al. ^35, 36^ and Fernández Rubio et al. ^37^ highlighted the crucial role of the cingulate gyrus, hippocampus, insula, inferior temporal cortex, and frontal operculum for the recognition of auditory sequences. The replication of previous findings encouraged us to further investigate the relationship between brain activity underlying auditory sequence recognition and individual WM skills.

Overall, this study showed a series of positive correlations between brain activity and WM abilities, suggesting that memory subsystems are coherently connected to each other. This is particularly interesting since the recognition task employed in the study used musical stimuli, while the WM measure was based on numbers. This link between different subsystems of memory is in line with previous research. As previously mentioned, the nature of the interactions between subsystems may be cooperative or competitive ^32, 33^ and is essential to perform memory tasks efficiently ^31^. Furthermore, different brain areas are involved depending on the memory process that is emphasized (declarative versus non-declarative) ^32^. Finally, White and McDonald’s ^34^ study localized multiple parallel memory subsystems in the rat’s hippocampus, caudate-putamen and amygdala, and proposed that these subsystems share information during learning, but represent its features differently.

Of particular interest in this study are the brain areas that were connected to WM. The activity recorded in the medial cingulate gyrus presented a sustained positive correlation with WM scores. This is coherent with previous studies linking cingulate gyrus’ activity to memory and musical tasks. As mentioned earlier, in the auditory domain, the cingulate played a crucial role in auditory sequence encoding ^39^ and recognition ^35-37^. Moreover, a recent meta-analysis revealed that the cingulate gyrus is central for general music processing and particularly for sound imagination ^40^. Beyond the auditory system, the cingulate gyrus has been reported in memory studies employing visual or abstract information. For instance, it has been suggested that diverse parts of the cingulate gyrus are differently involved in memory processes. According to this view, the anterior part of the cingulate is primarily connected to the orbitofrontal cortex and handles abstract, reward outcomes, while the posterior cingulate is integrated within the hippocampal and occipital systems and therefore highly relevant for memory processing of visual stimuli ^41, 42^. Similarly, in a recent fMRI study, Di and colleagues ^43^ showed that the anterior cingulate gyrus was functionally connected to the middle frontal gyrus and superior parietal lobule during a demanding, WM task. Conversely, this connectivity was reduced in resting state, suggesting the relevance of the cingulate gyrus during memory tasks.

Other brain structures correlated with WM abilities were the inferior temporal and temporal-fusiform gyri and the postcentral gyrus. This result is of great interest because these brain structures did not play a major role in the recognition of auditory sequences. Indeed, while the cingulate gyrus was largely active, we previously observed a relatively small contribution of inferior temporal and postcentral gyri to auditory sequence recognition ^35-37^. Moreover, the temporal-fusiform gyrus has not been previously associated to auditory recognition processes. This suggests that individuals with higher WM abilities recruited a larger brain network during recognition of auditory sequences, which may provide an advantage for auditory recognition. However, since there were no significant differences in the behavioral performance of the recognition task, future studies are called to better understand whether and how this recruitment of additional brain areas is beneficial for individuals with high WM capacity.

Previous literature has shown the involvement of inferior temporal and temporal-fusiform gyri and postcentral gyrus in visual memory tasks. In the past decades, the inferior temporal cortex has been widely connected to visual perception and memory in both humans and monkeys ^44^. Specifically, several studies demonstrated the involvement of the inferior temporal cortex in representational memory and recognition of complex visual patterns ^44, 45^. More recently, Costers and colleagues ^46^ reported the involvement of left and right inferior temporal and parahippocampal gyri in a multi-item WM task. Activity in the inferior temporal gyrus has been repeatedly observed in visual memory tasks, while its involvement in the auditory domain is less established. Importantly, here we revealed that the inferior temporal cortex plays a significant role in auditory recognition, at least in individuals with superior WM skills.

The fusiform gyrus has been historically connected to recognition in the visual domain, especially in relation to faces ^47-50^. However, recent studies demonstrated its involvement in the recognition and processing of more general visual stimuli, such as letters ^51^, and when performing elaborated associative learning tasks ^52^.

The postcentral gyrus is a brain area mainly associated to motor control ^53, 54^, yet evidence points to its contribution to memory processes. For instance, in a visual encoding task, a vast network of brain areas was active, including the postcentral gyrus ^55^. Similarly, in a recognition task of short sentences, supramarginal and postcentral gyrus activity was reported ^56^. Another study demonstrated the involvement of the postcentral gyrus in a WM and especially in a visual attention task ^57^. Notably, similar to the inferior temporal gyrus, previous literature reported activation of the postcentral gyrus mainly in relation to visual memory, while this study showed its involvement during recognition of auditory temporal sequences.

In conclusion, we discovered a positive correlation between individual WM abilities and brain activity underlying recognition of memorized auditory sequences, increasing our knowledge on the relationships between different memory subsystems. Future studies are encouraged to replicate our results and expand them by investigating the relationship between the brain mechanisms underlying recognition of temporal sequences and the brain processes associated to WM tasks.

## Materials and methods

### Data and code availability

The codes are available on GitHub (https://github.com/leonardob92/LBPD-1.0.git). The anonymized neuroimaging data from the experiment will be made available upon reasonable request.

### Participants

We recruited 71 participants (38 males and 33 females) who took part in the experiment on a voluntary basis. They were aged 18 to 42 years old (mean age: 25 ± 4.10 years). All participants were healthy and had normal hearing. Participants came from Western countries and had homogenous educational and socioeconomic backgrounds. Before starting the experimental procedures, participants gave their informed consent.

This study was a part of a larger project focused on brain dynamics underlying encoding and recognition of musical patterns. This project produced several studies ^35-37, 39^. In the current work, we used the brain activity data underlying recognition of musical patterns that was previously reported in Bonetti et al. ^35, 36^ and Fernández Rubio et al. ^37^ The project was approved by the Ethics Committee of the Central Denmark Region (De Videnskabsetiske Komitéer for Region Midtjylland, Ref 1-10-72-411-17). Moreover, the experimental procedures complied with the Declaration of Helsinki – Ethical Principles for Medical Research.

### Experimental stimuli and design

The study aimed at investigating the relationship between brain activity during a memory recognition task and working memory (WM) abilities (**Figure 1**).

The brain activity was measured using magnetoencephalography (MEG) while participants performed an old/new auditory recognition task. The task consisted of an encoding phase during which participants memorized a musical piece, and a recognition phase in which they recognized excerpts from the piece. In the encoding phase, participants were exposed to four repetitions of a full musical piece and were asked to memorize it as much as they could. The musical piece lasted for approximately 2.5 minutes. The total duration of the learning phase was approximately 10 minutes. For the recognition phase, 40 short excerpts (5-tone musical sequences, 1250 ms of duration in total) were extracted from the musical piece and 40 novel musical sequences were created. The resulting 80 sequences were presented in a randomized order. For each of them, participants were instructed to state whether the sequence was extracted from the musical piece they previously learned (memorized sequence) or whether it was a new sequence (novel sequence). To prevent from potential confounds, memorized and novel sequences were matched among several variables, including rhythm, timbre, volume, meter, tempo, number and duration of musical tones, tonality, information content (*IC*) and entropy (*H*).

This task was conducted independently for three musical pieces composed in different musical tonalities, with the aim of collecting a copious amount of data and increase the reliability of our findings. The three musical pieces were the right-hand part of J. S. Bach’s Prelude No. 1 in C minor BWV 847 (hereafter referred to as the “minor prelude”), the right-hand part of J. S. Bach’s Prelude No. 1 in C major BWV 846 (hereafter referred to as the “major prelude”), and an atonal version of the “major prelude” (hereafter referred to as the “atonal prelude”). All the pieces had the same duration. The atonal piece was composed by LB following a systematic change of pitch of the tones of the major prelude. Additional details on this procedure can be found in Fernández Rubio et al. ^37^

MIDI versions of the three pieces used in the encoding phase and the musical sequences used in the recognition phase were created using using Finale (MakeMusic, Boulder, CO) and presented to the participants through Presentation software (Neurobehavioural Systems, Berkeley, CA).

The WM abilities were assessed with the Wechsler Adult Intelligence Scale IV (WAIS-IV) ^38^, one of the most widely used tests to assess cognitive abilities. WAIS-IV comprises four main indices: Working Memory, Verbal Comprehension, Perceptual Reasoning, and Processing Speed. In this study, we used the two primary subtests of the Working Memory index: Digit Span and Arithmetic. In the Digit Span subtest, participants are required to repeat sequences of numbers either in the same order, backwards, or in ascending order, immediately after hearing them. In the Arithmetic subtest, participants have to solve mathematical problems without using any external aids (e.g., calculator, pen, etc.). These tests were performed outside the scanner.

### Data acquisition

The MEG data was recorded in a magnetically shielded room located at the Aarhus University Hospital (Denmark) with an Elekta Neuromag TRIUX MEG scanner equipped with 306 channels (Elekta Neuromag, Helsinki, Finland). The data was collected at a sampling rate of 1000 Hz with an analogue filtering of 0.1 – 330 Hz. Before starting the experiment, we recorded the participants’ headshape and position of four Head Position Indicator (HPI) coils with respect to three anatomical landmarks (nasion and left and right preauricular points) using a 3D digitizer (Polhemus Fastrak, Colchester, VT, USA). We used this information in a later stage of the analysis pipeline to co-register the MEG data with the MRI anatomical images. During the MEG experiment, the HPI coils recorded the continuous head localization, which was subsequently used to compensate for participants’ movement inside the MEG scanner. Moreover, two sets of bipolar electrodes were employed to record cardiac rhythm and eye movements. These were later used to remove electrooculography (EOG) and electrocardiography (ECG) artifacts.

The MRI scans were acquired on a CE-approved 3T Siemens MR-scanner at Aarhus University Hospital (Denmark). We recorded a structural T1 with a spatial resolution of 1.0 × 1.0 × 1.0 mm and the following sequence parameters: echo time (TE) = 2.96 ms, repetition time (TR) = 5000 ms, bandwidth = 240 Hz/Px, reconstructed matrix size = 256 × 256.

The MEG and MRI recordings were acquired in two separate sessions.

### Data preprocessing

The raw MEG sensor data (204 planar gradiometers and 102 magnetometers) was preprocessed by MaxFilter ^58^ in order to suppress external artifacts interfering with the magnetic field produced by the brain activity. Using MaxFilter, the data was also corrected for head motion and downsampled to 250 Hz. We then converted the data into Statistical Parametric Mapping (SPM) ^59^ format and further analyzed it in MATLAB (MathWorks, Natick, MA, USA) using the Oxford Centre for Human Brain Activity (OHBA) Software Library (OSL, https://ohba-analysis.github.io/osl-docs/), a freely available software that builds upon Fieldtrip ^60^, FSL ^61^, and SPM toolboxes, and in-house-built functions. We applied a notch filter to the data (48 – 52 Hz) to correct for inferences of the electric current. The signal was further downsampled to 150 Hz and the continuous MEG data was visually inspected to control for artifacts. To remove EOG and ECG components, we computed independent component analysis (ICA), isolated and discarded the components that picked up the EOG and ECG activity and reconstructed the signal with the remaining components. We then bandpass-filtered the data in the 0.1 – 1 Hz band, since we had previously shown ^35-37^ that activity in this slow frequency is mainly associated with the recognition of musical sequences. The data was subsequently epoched into 80 trials (40 memorized and 40 novel musical sequences), independently for the recognition of the three musical preludes. Then, we merged the three datasets, obtaining 240 trials (120 memorized and 120 novel musical sequences) without differentiating between the three musical preludes. Here, each trial lasted 3500 ms (3400 ms plus 100 ms of baseline time) and further analyses were performed on correctly identified trials only.

### Source reconstruction

After computing the preprocessing of the data, we estimated the brain sources which generated the signal recorded by the MEG. This procedure was carried out by designing a forward model and computing the inverse solution using beamforming algorithms) **Figure 1** shows an illustration of the source reconstruction pipeline.

Fist, using the information collected with the 3D digitizer, the MEG data and the individual T1-weighted images were co-registered, independently for each participant. We used the MNI152-T1 standard template with 8-mm spatial resolution in the case of four participants whose individual anatomical scans were not available.

Second, we computed a single shell forward model using an 8-mm grid. This theoretical head model considers each brain source as an active dipole and calculates how a unitary strength of such dipoles would be reflected over the MEG sensors ^62^. Then, we used a beamforming algorithm as inverse model. This is one of the most used algorithms for reconstructing the brain sources from MEG channels’ data. It consists of employing a different set of weights based on the forward model and the covariance between the MEG channels. Afterwards, these weights are sequentially applied to the source locations (dipoles) for computing the contribution of each source to the activity recorded by the MEG channels, independently for each time point ^63-65^.

### Brain activity underlying recognition of previously memorized versus novel musical sequences

Before evaluating the relationship between WM abilities and brain activity underlying musical sequence recognition, which was the main aim of the current work, we wished to replicate the established finding ^35-37^ that recognition of previously memorized versus novel auditory sequences is associated to a stronger activation of a widespread network of brain areas.

Thus, we first sub-averaged the brain data in five time-windows corresponding to the duration of the five tones of the musical sequences (0 – 250 ms, 251 – 500 ms, 501 – 750 ms, 751 – 1000 ms, 1001 – 1250 ms). Second, independently for the five time-windows, we computed one t-test for each brain source, contrasting the brain activity underlying recognition of previously memorized versus novel musical sequences. Third, we corrected for multiple comparisons using cluster-based Monte-Carlo simulations (MCS).

Cluster-based MCS returned the spatial clusters of brain sources that exhibited a significantly different activity between our two experimental conditions (α = .001). Then, the significant brain voxels emerged from the previous t-tests were shuffled in space and the maximum cluster size was measured. Repeating this procedure for each of the 1000 permutations used in the MCS analysis, we built a reference distribution of the maximum cluster sizes computed in the permuted data. Then, the original cluster sizes were compared to the reference distribution and were considered significant only if their size was bigger than the 95% of the maximum cluster sizes of the permuted data.

### WM abilities and brain activity underlying recognition of musical sequences

Before computing neural data analyses, we inspected whether there was a relationship between recognition accuracy and WM skills. To this aim, we computed a Pearson’s correlation between the individual WM scores (from WAIS-IV) and the number of correctly recognized auditory sequences in the MEG task.

To determine the relationship between WM abilities and brain activity underlying recognition of musical sequences, we computed Pearson’s correlations between participants’ WM scores and each of the reconstructed brain sources. We corrected for multiple comparisons using cluster-based MCS analogous to the ones described in the previous subsection. This procedure was computed independently for five-time windows that corresponded to the duration of the five tones of the musical sequences (0 – 250 ms, 251 – 500 ms, 501 – 750 ms, 751 – 1000 ms, 1001 – 1250 ms). Cluster-based MCS returned the spatial clusters of active brain sources during recognition of musical sequences that significantly correlated (α = .05) with the participants’ WM abilities. For each of the five MCS, the data was sub-averaged in the correspondent time window (as reported above) and the brain activity underlying recognition of novel sequences was subtracted from the brain activity underlying recognition of memorized sequences. In this way, we correlated the WM scores with the brain activity that was associated to the recognition of the sole memorized sequences. Then, the significant brain voxels emerged from the previous correlations were shuffled in space and the maximum cluster size was measured. Repeating this procedure for each of the 1000 permutations used in the MCS analysis, we built a reference distribution of the maximum cluster sizes computed in the permuted data. Then, the original cluster sizes were compared to the reference distribution and were considered significant only if their size was bigger than the 95% of the maximum cluster sizes of the permuted data.

## Acknowledgements

The Center for Music in the Brain (MIB) is funded by the Danish National Research Foundation (project number DNRF117).

LB is supported by Carlsberg Foundation (CF20-0239), Center for Music in the Brain, Linacre College of the University of Oxford, and Society for Education and Music Psychology (SEMPRE’s 50th Anniversary Awards Scheme).

MLK is supported by Center for Music in the Brain and Centre for Eudaimonia and Human Flourishing funded by the Pettit and Carlsberg Foundations.

We thank Giulia Donati, Riccardo Proietti, Giulio Carraturo, Mick Holt, and Holger Friis for their assistance in the neuroscientific experiment. We also thank psychologist Tina Birgitte Wisbech Carstensen for her help with the administration of psychological tests and questionnaires.

Finally, we thank the Fundación Mutua Madrileña for the economic support provided to the author Gemma Fernández Rubio and the University of Bologna for the economic support provided to student assistants Giulia Donati, Riccardo Proietti, and Giulio Carraturo.

## Data availability

The codes are available on GitHub (https://github.com/leonardob92/LBPD-1.0.git). The anonymized neuroimaging data from the experiment can be made available upon reasonable request.

## Author contributions

LB, GFR, MLK, FC, and PV conceived the hypotheses and designed the study. LB, GFR, and FC performed pre-processing and statistical analysis. LB, MLK, and PV provided essential help to interpret and frame the results within the neuroscientific literature. LB, GFR, and FC wrote the first draft of the manuscript and prepared the figures. All the authors contributed to and approved the final version of the manuscript.

## SUPPLEMENTARY MATERIALS

Supplementary materials related to this study are organized as supplementary figures and tables. Due to their large size, supplementary tables have been reported in Excel files that can be found at the following link: https://drive.google.com/drive/folders/1z3S7BTV7t5jfko6XDYprbdi84oJwgE9s?usp=sharing

## SUPPLEMENTARY TABLES

***Table ST1. Significant clusters of activity for MEG source data (memorized versus novel sequences)***.

Significant clusters of activity estimated from the contrasts between the brain activity (in 0.1 – 1 Hz) underlying memorized and novel musical sequences. The table depicts the contrast for each of the tones comprising the musical sequences, along with the brain regions, hemispheres, and *t*-values for each voxel.

***Table ST2. Significant clusters emerged from the correlation between WM abilities and MEG brain data underlying recognition of previously memorized musical sequences***.

Significant clusters of activity estimated from the correlations between WM abilities and the brain activity (in 0.1 – 1 Hz) underlying previously memorized musical sequences. The table depicts the correlation for each of the tones comprising the musical sequences, along with the brain regions, hemispheres, and *r*-values for each voxel.

